# BEES: Bayesian Ensemble Estimation from SAS

**DOI:** 10.1101/400168

**Authors:** Samuel Bowerman, Joseph E. Curtis, Joseph Clayton, Emre H. Brookes, Jeff Wereszczynski

**Affiliations:** Department of Physics and the Center for Molecular Study of Condensed Soft Matter, Illinois Institute of Technology, Chicago, IL 60616; Current address: Department of Biochemistry and Howard Hughes Medical Institute, University of Colorado Boulder, Boulder, CO 80309; NIST Center for Neutron Research, National Institute of Standards and Technology, Gaithersburg, MD 20899; University of Texas Health Science Center, San Antonio, TX 78229

## Abstract

Many biomolecular complexes exist in a flexible ensemble of states in solution which are necessary to perform their biological function. Small angle scattering (SAS) measurements are a popular method for characterizing these flexible molecules due to their relative ease of use and ability to simultaneously probe the full ensemble of states. However, SAS data is typically low-dimensional and difficult to interpret without the assistance of additional structural models. In theory, experimental SAS curves can be reconstituted from a linear combination of theoretical models, although this procedure carries significant risk of overfitting the inherently low-dimensional SAS data. Previously, we developed a Bayesian-based method for fitting ensembles of model structures to experimental SAS data that rigorously avoids overfitting. However, we have found that these methods can be difficult to incorporate into typical SAS modeling workflows, especially for users that are not experts in computational modeling. To this end, we present the “Bayesian Ensemble Estimation from SAS” (BEES) program. Two forks of BEES are available, the primary one existing as module for the SASSIE webserver and a developmental version that is a standalone python program. BEES allows users to exhaustively sample ensemble models constructed from a library of theoretical states and to interactively analyze and compare each model’s performance. The fitting routine also allows for secondary data sets to be supplied, thereby simultaneously fitting models to both SAS data as well as orthogonal information. The flexible ensemble of K63-linked ubiquitin trimers is presented as an example of BEES’ capabilities.

## 2 Introduction

Biological molecules rely heavily on their conformational dynamics to conduct their cellular function, and the characterization of these flexible ensembles of states remains a key challenge in modern biophysics^1^. As a result, many different experimental and computational techniques have been developed to probe and model configurational ensembles. Of these, small angle scattering (SAS) measurements are an increasingly popular technique due to their relative ease of use and ability to simultaneously probe the full solution ensemble^2,3^. Moreover, SAS measurements are able to probe systems at room temperature, free from packing forces induced by the lattice and cryogenic effects of crystallography, and they can measure the solution of states in both equilibrium ensembles and time-dependent processes^4^, such as protein and RNA folding^5,6^, or the allosteric coupling of enzymatic activity and large-scale domain movement^7,8^. However, the low-dimensional nature of SAS data can often cause the interpretation of scattering profiles to be relatively difficult, and reconstituting a three-dimensional molecular structure solely from scattering curves can often be misleading, as multiple reconstitutions of varying shapes may result from the same scattering profile.

In contrast, model structures can also be identified from all-atom or coarse-grained simulations, and their calculated scattering profiles can be compared against empirical curves^9–12^. Since SAS profiles are measurements of the full solution ensemble and therefore may not be fully described by a single structural state, these in *silico* profiles can also serve as a basis set to construct an ensemble model through a linear combination of states^13–16^. While this ensemble reconstitution approach is conceptually straightforward, in practice it can be quite difficult to identify the “best” ensemble model. For instance, it is not known *a priori* what the number of underlying states should be in the ensemble. It is also possible for ensemble models to overfit experimental data through the inclusion of too many underlying populations. Furthermore, altogether different combinations of states may yield similarly performing models, in respect to their goodness-of-fit values.

For these reasons, a Bayesian-based approach has many advantages over more traditional methods. For instance, Markov Chain Monte Carlo posterior sampling methods will not only estimate model parameters but will also allow for the direct assessment of their errors^17^. Moreover, Bayesian formalism allows for the comparison of a population of models as a solution to parameterization, rather than only identifying a single set of parameters^18–21^. This is exceptionally useful for SAS modeling, where information regarding the model is underdetermined. However, the ability to construct a large population of solutions can also be a disadvantage, as both the computational resources to construct a complete array of model parameters, as well as tools for comparing models, can be daunting for many systems.

To this end, we previously developed an iterative Bayesian method to use small angle scattering (SAS) profiles, either of x-rays (SAXS) or of neutrons (SANS), to re-weight the population of states from simulated models. This approach, which is an extension of the BSS-SAXS technique^13^, compares solution ensembles of a variety of sub-ensembles from a combination of potential scattering states. Originally, we used this method to fit ensembles of covalently linked ubiquitin trimers, and we observed that the algorithm could produce ensemble models that robustly resisted overfitting^22^.

Here, we present an update to this method as an open source program called “Bayesian Ensemble Estimation from SAS” (BEES, henceforth). Two versions of this code have been developed. The primary version is an open-access module on the SASSIE-web server (http://sassie-web.chem.utk.edu/sassie2/), which provides a graphical user interface for controlling the module^23,24^. The BEES-SASSIE module is designed for users that are both new and experienced in biophysical modeling, and, through SASSIE, it provides access to the computational resources required to calculate and analyze large combinations of states. The second, developmental, version is a stand-alone python code that is designed to be run from the command line, and is intended for experienced computational scientists. We also provide two example use cases, one in which we fit profiles of K63-linked ubiquitin trimers to SAXS data alone and another in which we add a second data set to the fitting procedure.

## 3 Methods

### 3.1 BEES Algorithm

The BEES algorithm is designed to find the theoretical solution ensemble that uses the fewest number of populations to accurately describes the experimental data. This algorithm is briefly presented here (Fig 1), but further details can be found in the supplemental text and elsewhere^22^. In short, experimental data are gathered and post-processed prior to using the BEES module. For example, users may wish to screen their data for low-q beam smearing effects or to extrapolate their scattering profile to I(0). A collection of theoretical profiles for candidate solution states are also input to BEES, which can be computed by standalone programs such as Crysol^25^ or FoXS^26^, in SASSIE via the “SasCalc” module^27^, or from many other scattering prediction software ^28–31^.

**Figure 1:**
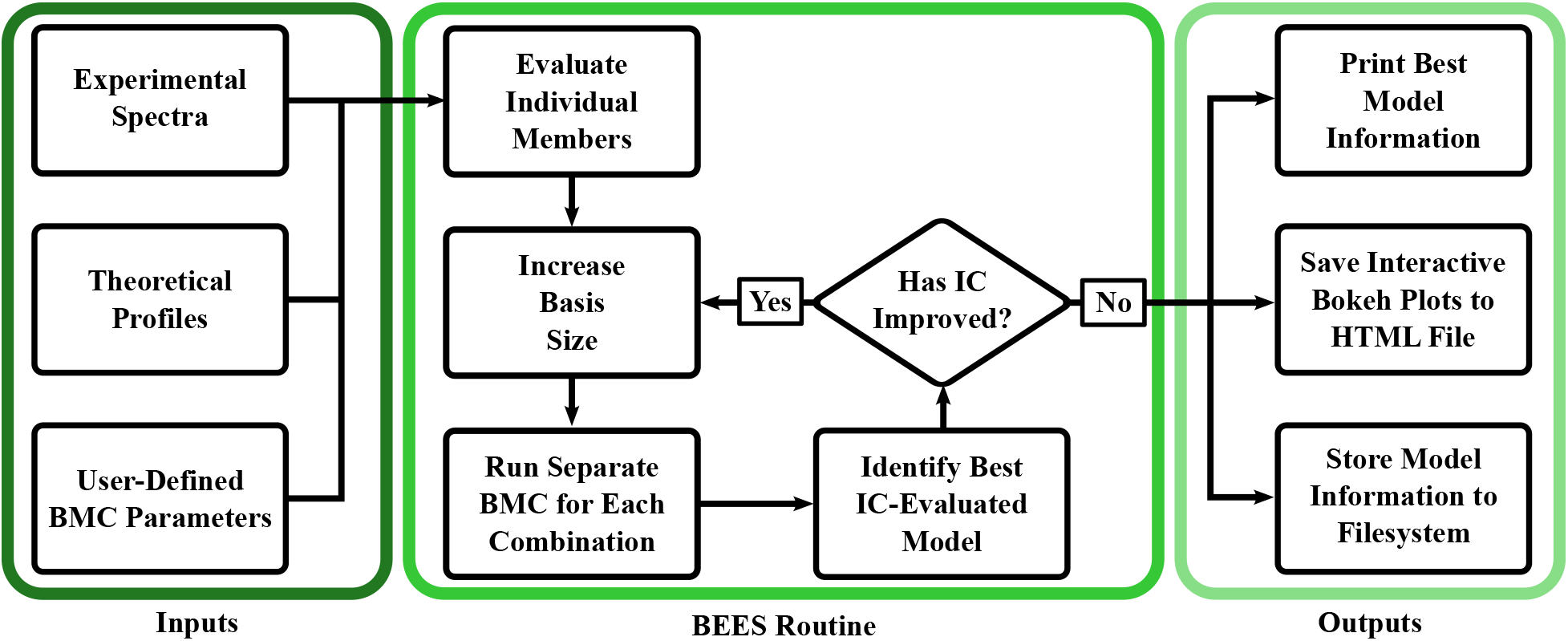
Workflow schematic of the BEES routine. Users supply empirical data and the collection of theoretical profiles for potential ensemble members, as well as set several parameters associated with the Bayesian Monte Carlo (BMC) parameter search. After the performance of each individual theoretical state is evaluated, ensemble populations are fit by BMC routines conducted iteratively on increasing sized sub-ensembles, until the addition of another member population does not improve the IC value and overfitting is observed. Alternatively, users can bypass the IC-comparison step to compare all possible combinations of states. The routine then relays information regarding the resulting models to the command terminal (stand-alone version) or GUI (SASSIE-web version) and further stores model information in several file locations for further review by users.

Once initiated, the BEES routine first determines the goodness-of-fit values of each individual profile. It then identifies all possible sub-bases containing combinations of two theoretical profiles, and it conducts a Bayesian Monte Carlo routine on each combination to identify the population of states in each sub-basis. Each Monte Carlo routine is conducted according to user-defined parameters: number of independent Monte Carlo parameter fittings per sub-basis, number of iterations per Monte Carlo fitting, and amount of population change per iteration. Notably, the BEES likelihood function (*L*) includes the ability to simultaneously fit the scattering profiles and an auxiliary set of measurements:

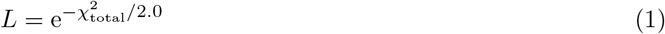

where the total model goodness-of-fit 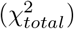 is the linear combination of the model scattering goodness-of-fit 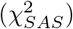 and the model goodness-of-fit to the auxiliary data set 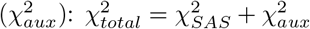.

Once the ensemble of states for each two-member sub-basis has been identified, the best two-member state is selected in accordance to the information criteria (IC) selected by the user, either the Akaike information criterion^32^ or the Bayesian Information Criterion^33^ (see Section 3.2 for more details). If the IC value of the best two-member state is worse than that of any single theoretical profile, then the module reports the best single profile as the most likely model. However, if the IC value of this two-member state is instead an improvement over all individual profiles, then the BEES module conducts the Bayesian Monte Carlo routine on every three-member sub-basis, and the best three-state IC value is similarly compared to the two-state ensemble. This iterative increase in sub-basis size and comparison of IC values is conducted until either the IC metric does not improve or every possible combination of states is considered. Alternatively, users also have the option to override the IC-comparison and force the construction of all combinations of subensembles. Once the desired number of models have been identified, the BEES module will also calculate each model’s “relative performance” metric to determine its likelihood over the best IC-identified model (Section 3.2)^34^:

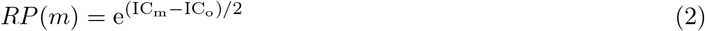

where *RP*(*m*) and *IC_m_* are the the relative performance and IC values of model *m*, and *IC_o_* is the minimum IC value of all observed models. The relative performance metric is more commonly known as the relative likelihood of a model. Here, we opt for the changed nomenclature to assist non-experts in the interpretation of the metric, as well as to avoid confusion with the likelihood function used by the Bayesian Monte Carlo fitting routine. While the relative performance provides a quantitative result, it is admittedly an approximation of the more rigorous Bayes Factor^33,35^. As such, it is intended to be interpreted loosely and to assist the user in applying their intuition toward the performance of alternative ensembles to the best identified one.

Once the best model has been identified, BEES outputs information regarding ensemble members of the IC-identified model, its model population weights, goodness-of-fit information for the full ensemble model and each individual, and the IC value of the model. Beyond the best identified model, information regarding every model identified for each sub-basis is also saved. Plots of the model ensemble fit to the experimental data, along with the associated residual errors, are automatically created once the fitting routine is completed. These plots are included in a multi-tab HTML page that which provides graphical and table presentations to allow users the ability to compare different models and performances.

### 3.2 Comparing Model Perfomances with Information Criteria

The rigorous comparison of theoretical ensembles to experimental data requires creating models that are rich enough to describe the underlying physical structures that generated the data while simultaneously avoiding overfitting. Biomolecules exist in an ensemble of conformations in solution, therefore an ensemble of theoretical structures is typically required to interpret SAS data. However, it is imperative that the final model does not achieve a strong goodness-of-fit value through inclusion of an arbitrary number of parameters (here, the number of scattering profiles). As a result, the true “best model” must be a balance between optimizing the goodness-of-fit metric and minimizing the number of underlying scattering states. To this end, the BEES module utilizes “Information Criterion” (IC) in order to penalize model goodness-of-fit values according to their ensemble size. Users have the option to use one of two different IC metrics during fitting — the Akaike Information Criterion (AIC) or the Bayesian Information Criterion (BIC)^32,33^:

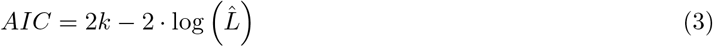

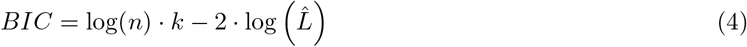

Here, *k* is the number of model parameters (number of scattering states), 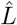 is the maximum observed likelihood value during the Bayesian Monte Carlo parameter fitting, and n is the number of points in the experimental data set.

Both the BIC and AIC have forms that reward models with improved experimental fits (higher values of 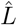) and penalize those with more parameters (higher values of *k*). The BIC is closely related to the AIC; however, it is derived from Bayesian principles rather than the frequentist foundation of the AIC. In both metrics, smaller values are indicative of better model performance, with the defining separation between them being the strength of the penalty term. In the AIC, the penalty is always double the number of states, whereas the BIC penalty will become increasingly larger for a larger number of data points. In reality, both metrics are an approximate way to identify the true model, and the AIC may be more prone to false positive estimations (including too many states), while the BIC metric may be more prone to false negatives (rejecting too many states), depending on the number of experimental data points. However, it is often possible that both metrics converge upon the same solution, as is the case with the K63 example presented here.

The model with the minimum IC value can be interpreted as the most likely, best performing, model. While it may be tempting to accept this model and reject all others, Bayesian principles dictate that there is a possibility that one of these other models might actually be more accurate to the true nature of the system, even though each one possesses a weaker IC value. The probability that a model is, in fact, a better assessment of the data can be calculated by comparing the model IC values to the lowest IC value, as previously stated (Eqn 2)^34^.

Because the BIC and AIC apply different penalties to the number of states, they may also produce different relative performance values for the same set of models. Depending on the number of independent data points, the BIC will produce relative performance values for competing models that are either closer to (n ≤ 7) or further from (n ≥ 8) the performance of the model with the lowest BIC. That is, if the number of independent data points is seven or fewer, then more models will have a relative performance closer to 1.0 than if evaluated by AIC. On the other hand, if the number of observed data points is greater than eight, then more models will have relative performances closer to 0.0 if they are evaluated by the BIC in place of the AIC. In the end, the choice of BIC vs AIC evaluation is up to the user, and it may sometimes be appropriate to use both to determine upper and lower bounds for relative model performances.

## 4 Results

Here, we describe a sample usage of BEES and its resulting data. The necessary data files for this test set are included in the Supporting Information. Users can thereby re-create the analyses presented here by unpacking the archive locally and uploading the relevant files for each case to the BEES module in SASSIE-web, or by following the shell scripts provided alongside the stand-alone version (https://github.com/WereszczynskiGroup/BEES/tree/master/examples). In the first example, we model the populations of states of K63-linked ubiquitin trimers using clusters identified from accelerated molecular dynamics trajectories^22^. In the second example, we showcase the effects of simultaneously fitting the SAS spectra and an auxiliary data set by including simulated measurements of an inter-domain distance and angle.

### 4.1 Building Ensembles of SAS Data

BEES requires the user to supply the experimental scattering curve along with theoretical scattering curves for candidate structures. In addition to providing this data, users must also define the D_max_ of the molecule, which can be determined from the experimental profile using pre-existing software^36^. Here, a D_max_ of 83.6 Å was determined using the Shanum program of the ATSAS package^37^. Furthermore, five Monte Carlo walkers were used for each sub-basis ensemble, and each walker was conducted for 10,000 iterations. The first 1,000 iterations were neglected when determining the model populations so as to remove any influence of the randomly selected initial values from the final result. Parallel processing can also be used (here, six processors were used), but using multiple processors will only enhance the speed of the calculation and has no effect on the final result (see Supporting Information for more information). In addition, the full array of sub-ensembles has been calculated to display the depth of analysis available. In this example, truncation of the algorithm via the IC parameter would save a significant amount of computational time without effecting the best IC-identified model; however, models with lower χ^2^_free_ would not have been observed. At the conclusion of the BEES routine, the best identified model is reported (Fig 2A), and an interactive plot interface is created (Fig 2B).

**Figure 2:**
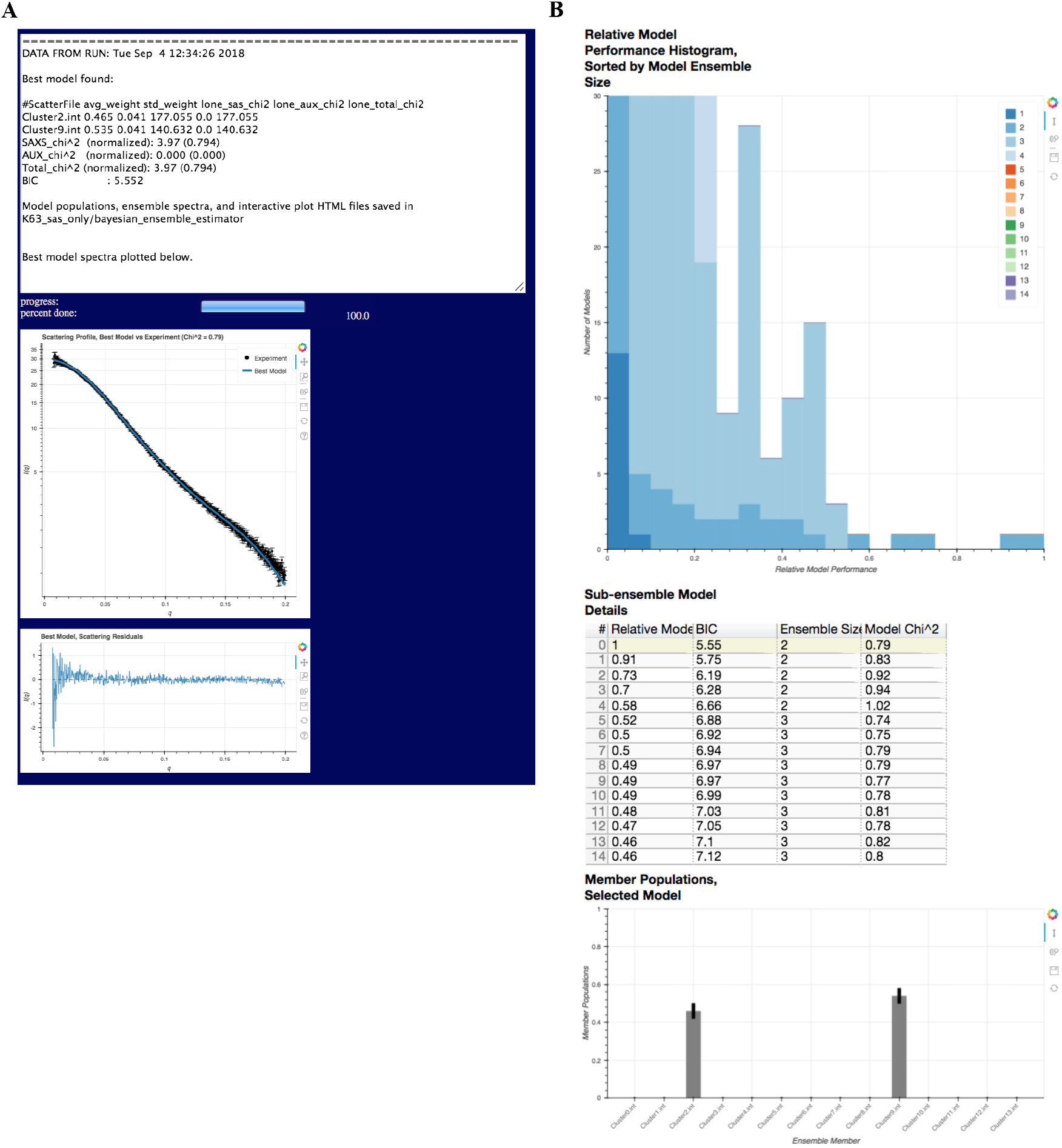
(a) Example of output from BEES program, as used on SASSIE-web GUI. (top) Text output displaying the contributing populations of the best IC-identified ensemble and the associated error in population estimates, as well as goodness-of-fit for each member. Total model goodness-of-fit and IC value are also printed by the module. (middle) Ensemble scattering profile of the best identified model, shown in blue, fit to the experimental spectrum, shown in black. (bottom) Residual errors of the best model against the experiment. (b) The third tab of the BEES-output HTML file (“Compare All Models”), which contains the relative performances histogram as well as a table of all the constructed ensemble models and their relative performance, ensemble size, selected IC metric, and goodness-of-fit values. Selecting a particular model in the table will also visualize the constituent populations on the bar graph below (best identified model selected here). The full interactive HTML file can be accessed by downloading the “K63_sas_only_plots.html” file from the example files contained within the Supporting Material. A similar file for the inclusion of auxiliary data can be found in “K63_with_aux.html”, also included in the Supporting Material example files.

In this example, the best model is a two-state solution that is approximately equal parts clusters 2 and 9. This model has a χ^2^_free_ of 0.79 and a BIC value of 5.55. While this is the best model according to BIC comparisons, roughly 50 models of varying sizes possess better χ^2^_free_ values, and the model with the best goodness-of-fit (χ^2^_free_ = 0.74) is a 4-member state comprised of clusters 2 (~45%), 4 (~22%), 10 (~15%), and 11 (~18%). This lowest χ^2^_free_ model has an IC value of 8.47, which yields a relative performance of 0.23 when compared to the IC-identified two-state model. As such, the improved χ^2^_free_ value of this model is unwarranted, as it is likely the result of overfitting by too many basis members. Indeed, inspection of the model performance histogram (Figure 2B, top) shows that the best performing models are largely two-state solutions, but some three-state solutions perform moderately well. Furthermore, many of the two- and three-state solutions are a significant improvement over each of the single-state models.

### 4.2 Building Ensembles with Auxiliary Data

Some users may desire to use BEES to build theoretical solution states by fitting solely to SAS data, and then use these states to predict measurements of future experiments. However, others may already possess such data and may prefer to create models that are consistent with both these measurements as well as the observed SAS profiles. For example, an experimenter may desire to simultaneously model both a scattering profile and a catalogue of NMR-derived distances. For the benefit of this class of users, we have included this functionality within BEES. To demonstrate how including such data might affect the modeling results, we discuss here an extension of the previous tri-ubquitin example in which we provide a simulated data set that contains the ensemble-averaged center-of-mass distance between distal monomers and the angle formed by the trimer arrangement (Fig 3). These data were created by taking the ensemble-averaged measures of the best model from the previous example with the inclusion of a Gaussian noise factor, resulting in a target distance of 53.0 ± 1.6 Å and a target angle of 117.7 ± 8.3°. Inputs to the BEES routine are identical to the previous example, with the exception of the auxiliary data set.

**Figure 3:**
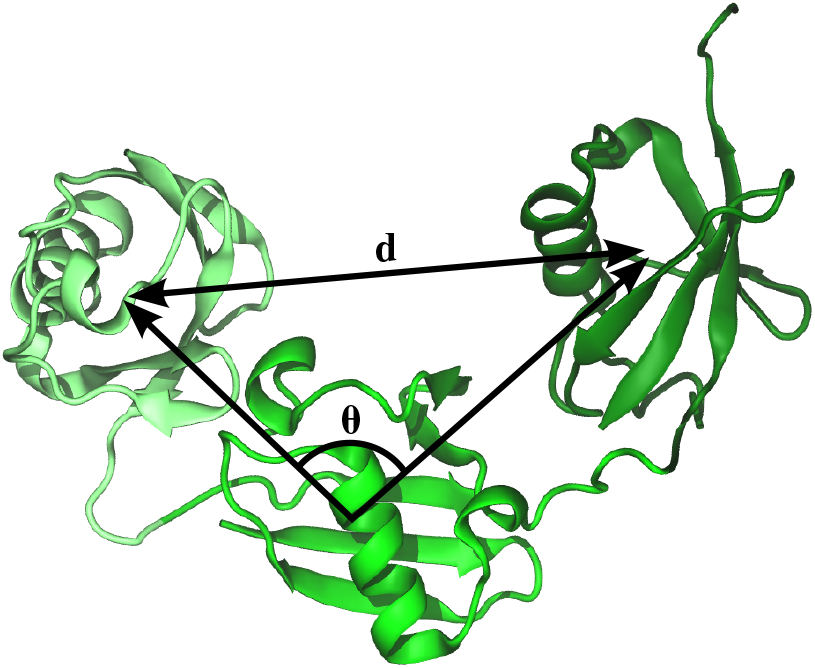
A visual representation of the two auxiliary measurements included in the second BEES routine. Both the distal monomer separation distance (d) and angle (*ϑ*) are measured in accordance to each monomer’s center-of-mass.

With the addition of the distance and angle measurements, we find a shift in the best IC-identified model. While still a two-state solution, the contributing members are now clusters 3 (43 ± 5%) and 4 (56 ± 5%). This model yields a χ^2^_total_ of 0.80, with a χ^2^_SAS_ of 0.96 and a χ^2^_aux_ of 0.38. As was the case in the last example, there are a plethora of models containing three or more members in which better goodness-of-fits are observed, and the best goodness-of-fit model is a mixture of clusters 2, 4, and 11 and has a χ^2^_total_ of 0.65. While this model is arguably a better fit to the data than the two-state ensemble of clusters 3 and 4, the IC value of this model is larger due to the addition of a third population. As such, this model is only the eighth most probable model, and possesses a relative performance of 0.63.

When we inspect the ten best ensembles, we once again find the best model from the previous example, which possesses a χ^2^_total_ of 0.81, a χ^2^_SAS_ of 0.81, and a χ^2^_aux_ of 0.83. Differences between the exact values of the χ^2^_SAS_ metric in this example and the previous example are a result of the random-sampling nature of the χ^2^_free_ metric, but these values are statistically indistinguishable. Similarly, the total goodness-of-fit in the clusters 3 and 4 ensemble is comparable to the ensemble containing clusters 2 and 9. As both models are two-state solutions, this results in very similar IC metrics and a relative performance value of 0.94, which suggests that neither model is significantly more accurate than the other. However, the 3+4 ensemble significantly outperforms the 2+9 ensemble in the context of the distance and angle measurements, while the 2+9 ensemble is a better fit to the the scattering curve.

## 5 Discussion

Here, we have presented the Bayesian Ensemble Estimation from SAS (BEES) program and highlighted its use with two example use cases. In the first example, we used the module to reweight states of K63-linked tri-ubiquitin that were obtained from accelerated molecular dynamics simulations. The BEES module identified a two-state solution as the model that best balanced the fit to experimental data with the fewest number of states. However, the analysis also found a plethora of models that had improved goodness-of-fits to the experimental scattering profile, but each of these models had more ensemble members than the two-state solution. The BEES module provides users with a convenient interface to both find and compare these other candidate ensembles with the IC-identified best state. This allows researchers the option to either rigorously trust the IC statistics to identify the most appropriate scattering model or to use the “ensemble of ensembles” constructed by the BEES module to guide their understanding of datasets separate from the fitting procedure.

The second use case discussed here demonstrated how BEES performs when simultaneously fitting populations to both SAXS and auxiliary data (here, simulated distance and angle measurements). In this example, the best identified model was still a two-state solution. However, a three-member ensemble was observed to have a better goodness-of-fit, but the improvement to χ^2^_total_ was not sufficient to also improve the IC parameter, yielding a relative performance of 0.63. Since the two-state solution has strong agreement with both measurements (χ^2^_free_, χ^2^_aux_ < 1.0), this relative performance value suggests that a conservative estimate for the solution ensemble would favor the two-state model over the χ^2^ three-state case. However, the performance is of high enough quality that this ensemble could also be considered as a solution for future measurements. In this way, we emphasize that the relative performance metric should aide the intuition of researchers, rather than completely replace it.

BEES seeks to identify the theoretical ensemble of states that uses the fewest number of populations to accurately describe the experimentally measured solution ensemble. In doing so, BEES is biased toward fitting the minimum amount of information contained within the experimental data, so as to avoid potential over-fitting. In contrast, other methods such as genetic algorithms and maximum entropy approaches will seek to use the full information of each scattering point^14,15,38^. While these methods may result in overfitting, BEES is also susceptible to under-fitting when utilizing SAXS data alone. As a result, the most accurate model to the true solution ensemble is likely one that is of a size between the smallest and largest ensembles identified by these methods. Furthermore, accurate use of any of these fitting methods is reliant on high-quality theoretical profiles; inaccurate theoretical states will likely lead to incorrect models. Therefore, users should be very careful when selecting scattering calculator programs and parameters, and special attention should be paid to accurately accounting for hydration layer effects^39^.

BEES can be used to construct ensemble models of scattering data from a library of candidate states, and the iterative algorithm of BEES quantitatively resists overfitting of the data from the addition of unnecessary populations. The program is available as a module on SASSIE (https://sassie-web.chem.utk.edu/sassie2/), as well as in a stand-alone form (https://github.com/WereszczynskiGroup/BEES). BEES is designed for use by both new and expert users of computational ensemble modeling, and the GUI-based module for the SASSIE-web platform provides structural and computational biophysicists with the resources necessary to construct molecular models in a Bayesian-based manner. Furthermore, BEES provides visual tools for quickly interpreting not only the quality of the best IC-identified model, but also for the full ensemble of sub-basis models available from the candidate populations. This feature allows users to inspect many different potential solutions and to compare their ability to model both SAS and auxiliary data sets. In this way, BEES serves the intuition of structural researchers in building ensembles of states for their systems of interest.

## 6 Author Contributions

SB and JW designed the BEES routine; SB and JC wrote the code of the BEES routine; SB, JEC, and EHB designed and wrote the SASSIE-web implementation of BEES; SB, JC, and JW analyzed the data sets; SB and JW wrote the first manuscript draft, and all authors contributed to editing of the manuscript.

## 7 Acknowledgements

The authors would like to thank Dr. Susan Krueger for valuable discussions in designing the plotting interface. EHB’s work is supported by National Science Foundation grant number OAC-1740097 and NIH grant GM120600. SB, JC and JW are supported by the National Institute of General Medical Sciences (NIGMS) of the National Institutes of Health under award number R35GM119647. The content is solely the responsibility of the authors and does not necessarily represent the official views of the National Institutes of Health. This work benefited from CCP-SAS software developed through a joint EPSRC (EP/K039121/1) and NSF (CHE-1265821) grant, as well as interactions and data collection at the Biophysics Collaborative Access Team, which is supported by NIGMS grant P41GM103622. This work used the Extreme Science and Engineering Discovery Environment (XSEDE), which is supported by National Science Foundation grant number ACI-1548562^40^.

Certain commercial equipment, instruments, or materials are identified in this paper to foster understanding. Such identification does not imply recommendation or endorsement by the National Institute of Standards and Technology, nor does it imply that the materials or equipment identified are necessarily the best available for the purpose.

