## Supplemental Information for "BEES: Bayesian Ensemble Estimation from SAS"

### S1 SASSIE-web Framework

The primary version of BEES is contained within the SASSIE-web server. SASSIE-web utilizes the GenApp framework to create a graphical user interface (GUI) and facilitate the use of several independent scientific applications into modular workflows[1,2]. These applications, including the BEES module, are written primarily in Python (v2.7.13) and utilize libraries contained within the Anaconda distribution. BEES, both in the SASSIE and stand-alone versions, makes use of the mpi4py (v2.0.0)[3], NumPy (v1.13.1)[4], and SciPy (v0.19.1)[5] packages to optimize performance. It also utilizes the Bokeh (v 0.12.16)[6] library to create interactive plots and html files to assist with the interpretation of the ensemble models identified by the BEES algorithm.

### S2 BEES Parallelization Scheme

Since the BEES algorithm identifies the best ensemble model as a sub-basis of the full list of supplied potential states, the required CPU time for a single BEES analysis relies mainly on two parameters: the total number of theoretical profiles and the sub-basis size at which over-fitting is observed. While the number of theoretical profiles can be reduced by conducting a clustering analysis or by screening the profiles in accordance to orthogonal information before being input to BEES, the number of states before overfitting is observed is not known *a priori* and may likely be affected by the signal-to-noise ratio of the experimental data. However, because the Monte Carlo reweighting procedure of each sub-ensemble is independent of the others, the algorithm benefits substantially from a trivial parallelization scheme. Therefore, BEES utilizes mpi4py to separate sub-ensembles at each basis-size iteration in an Open MPI-compatible manner[3,7]. In this way, the BEES module on SASSIE-web provides users that possess limited parallel resources with the opportunity to conduct ensemble re-weightings on large basis sizes ($n > 10$ members) in a reasonable amount of time ($\sim$2 hours on 6 processors for 14 candidates that overfit at three member states, $\sim$36 hours on 6 processors for calculation of the full array of models; detailed further below). Similarly, users of the command line version may take advantage of their local parallel computational resources to reduce the required computational time.

#### S2.1 Parallel Performance

The BEES algorithm suffers from two potential bottlenecks: the number of profiles in the user-supplied candidate pool and the ensemble size at which overfitting is observed. While knowledge of the latter cannot be determined *a priori*, benchmarks of the BEES parallel performance can help guide users in determining how much data reduction is necessary before running the BEES module. These benchmarks were performed under minimal server load, and therefore represent a best case scenario (i.e., a lower limit to the computational wall clock) for the BEES algorithm. Detailed further below, we find that users can calculate the full combination of models from 14 in $\sim$36 hours using 6 processors, and candidate pools of 22 members can be fit to three-member states (overfitting observed in four-member models) in the same timescale using 6 processors.

To perform these benchmarks, we constructed candidate states of varying sizes (n = 2, 6, 8, 10, 12, 20, 25) in addition to our original 14-member candidate pool. Each BEES routine was run using five Monte Carlo replicas, where each replica was conducted for 10,000 iterations. It can be seen that the total number of combinations scales exponentially with the number of candidate profiles (Figure S1), as does the required wall clock. Furthermore, the time required to build the models scales almost linearly with the number of processors: for n = 10, the required time was 6.4 hours on two processors, in comparison to the 2.2 hours required for six processors. From these data, we observe that users can calculate full combinations of states from candidate pools of 14 members in a reasonable ($\sim$36 hours) time frame when run on 6 processors.

In contrast to the previous discussion, many users may not desire to calculate the full combination of states, but rather find the single best model that avoids overfitting. In this case, the required computational cost may be drastically reduced, especially if overfitting is observed at a relatively small ($\leq 3$) ensemble size. For this reason, we have conducted a secondary benchmark of BEES routines containing 10, 14, 20, and 25 candidate profiles (Figure S2). Each BEES run utilized five Monte Carlo replicas of 10,000 iterations each on six processors. Using these benchmarks, we see that a three-member solution (overfitting observed at four states) from a candidate pool of 25 members can be found in ∼30 hours under ideal server load.

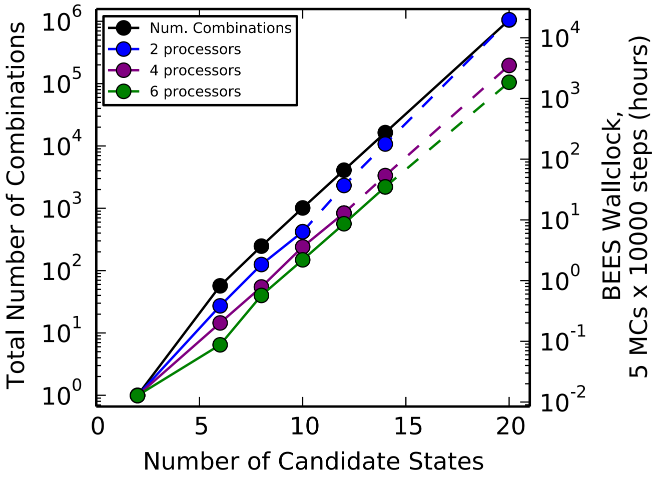


**Figure S1:** Performance of the BEES module with respect to candidate pool size when evaluating every possible combination of states. Benchmarks were determined for BEES routine of five Monte Carlo replicas that were run for 10,000 iterations each on two (blue), four (purple), and six (green) processors. Solid lines connect benchmarks that were explicitly measured, whereas dotted lines connect to points that are extrapolated from an exponential fitting of the measured benchmark data. From these data, even on six processors, the BEES module would require over 1,000 hours (∼42 days) to examine every possible combination of states from 20 candidate members.

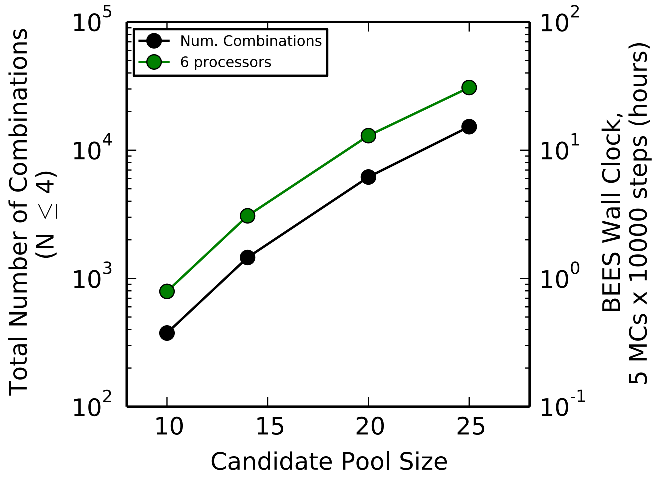


**Figure S2:** Performance of BEES module with respect to candidate pool size when the experimental data is best described by a three-state solution (overfitting observed at four-member solutions). Benchmarks were determined for a BEES routine that employed five Monte Carlo replicas that were run for 10,000 iterations on six processors. From the interpolation of this data, a three-state solution could be identified as the best model from a candidate pool of 25 members in ∼30 hours when run on six processors.

[6] Bokeh Development Team. Bokeh: Python library for interactive visualization, 2014.

[7] Edgar Gabriel, Graham E. Fagg, George Bosilca, Thara Angskun, Jack J. Dongarra, Jeffrey M. Squyres, Vishal Sahay, Prabhanjan Kambadur, Brian Barrett, Andrew Lumsdaine, Ralph H. Castain, David J. Daniel, Richard L. Graham, and Timothy S. Woodall. Open MPI: Goals, concept, and design of a next generation MPI implementation. In Proceedings, 11th European PVM/MPI Users' Group Meeting, pages 97–104, Budapest, Hungary, September 2004.
